# In the Body’s Eye: The Computational Anatomy of Interoceptive Inference

**DOI:** 10.1101/603928

**Authors:** Micah Allen, Andrew Levy, Thomas Parr, Karl J. Friston

## Abstract

A growing body of evidence highlights the intricate linkage of exteroceptive perception to the rhythmic activity of the visceral body. In parallel, interoceptive inference theories of emotion and self-consciousness are on the rise in cognitive science. However, thus far no formal theory has emerged to integrate these twin domains; instead most extant work is conceptual in nature. Here, we introduce a formal model of cardiac active inference, which explains how ascending cardiac signals entrain exteroceptive sensory perception and confidence. Through simulated psychophysics, we reproduce the defensive startle reflex and commonly reported effects linking the cardiac cycle to fear perception. We further show that simulated ‘interoceptive lesions’ blunt fear expectations, induce psychosomatic hallucinations, and exacerbate metacognitive biases. Through synthetic heart-rate variability analyses, we illustrate how the balance of arousal-priors and visceral prediction errors produces idiosyncratic patterns of physiological reactivity. Our model thus offers the possibility to computationally phenotype disordered brain-body interaction.

## Introduction

The enactive view of perception – implied by active vision and inference – suggests an intimate co-dependency between perception and the active sampling of our sensorium. In this work, we take the embodied view to its ultimate conclusion and consider perception as a function of the physical and physiological body we use to ‘measure’ the world. In particular, our focus is on the coupling – or interaction – between interoceptive and exteroceptive perception; namely, how bodily states and states of affairs beyond the body are inferred – and how inference about each domain affects the other. For example, does what we see depend upon our autonomic status and how does visual perceptual synthesis affect sympathetic or parasympathetic outflow? The body is, in essence, an ensemble of fluctuating systems with biorhythms nested at multiple timescales. How then do these physiological fluctuations interact with perceptual synthesis in the visual and auditory domains?

There is a rapidly growing body of evidence suggesting that bodily and autonomic states affect perceptual and metacognitive decisions (Allen et al., 2016b; Azevedo et al., 2017; Bonvallet and Bloch, 1961; Cohen et al., 1980; Garfinkel et al., 2014; Hauser et al., 2017b; Lacey and Lacey, 1978; Park et al., 2014; Salomon et al., 2016; Velden and Juris, 1975; Zelano et al., 2016). Much of this evidence emphasises the dynamic aspect of our physiology; usually assessed in terms of how psychophysics depends upon the phase of some physiological cycle. Most of the empirical evidence suggests that biorhythms gate or modulate the way that sensory evidence is accumulated during perception (Bonvallet et al., 1954; Bonvallet and Bloch, 1961; Karavaev et al., 2018; Varga and Heck, 2017). In the predictive coding literature, this is usually treated as fluctuating, context sensitive, changes in the precision of sensory sampling (e.g., the precision or gain of prediction errors). Clear examples of this include the fast waxing and waning of precision during active visual sampling. For example, saccadic suppression – during saccadic eye movements – alternates with attention to fixated visual information every 250 ms or so. This process of actively sampling the environment via ballistic saccade itself varies with the cardiac cycle (Galvez-Pol et al., 2018; Kunzendorf et al., 2019; Ohl et al., 2016). At still slower timescales, respiratory (Herrero et al., 2017; Tort et al., 2018b, 2018a; Zelano et al., 2016) are coupled to neuronal oscillations and behavior. In short, at probably every timescale there are systematic fluctuations in the precision or quality of sensory evidence that depend upon when we actually interrogate the world, in relation to the biorhythms of our sensory apparatus; namely, our body.

Our focus on the multimodal integration of interoceptive and exteroceptive domains is driven by the overwhelming evidence for interoception as a key modality in hedonics, arousal, emotion and selfhood (Allen and Friston, 2018; Apps and Tsakiris, 2014; Gallagher and Allen, 2018; Seth, 2013; Seth and Friston, 2016). This is generally treated under the rubric of interoceptive inference; namely, active inference in the interoceptive domain. There are several compelling formulations of interoceptive inference from the perspective of neurophysiology, neuroanatomy and, indeed, issues of consciousness in terms of minimal selfhood. However, much of this treatment rests upon a purely conceptual analysis – underpinned by some notion of active (Bayesian) inference about states of the world (including the body). In this work, we offer a more formal (mathematical) analysis that we hope will be a point of reference for both theoretical and empirical investigations.

In brief, we constructed a (minimal) active inference architecture to simulate embodied perception and concomitant arousal. Here, we focused on simulating interactions between the cardiac cycle and exteroceptive perception. In principle however, our simulation provides a computational proof-of-principle that can be expanded to understand brain-body coupling at any physiological or behavioral timescale. Using a Markov decision process formulation, we created a synthetic subject who exhibited physiological (cardio-acceleration) responses to arousing stimuli. Our agenda was twofold: first, to provide a sufficiency proof that – in at least one example – the interaction between interoception and exteroception emerges from the normative (formal) principles of active inference. Furthermore, having an *in silico* subject at hand, means that we can simulate the effects of various disconnections and pathophysiology. For example, we can examine the effect of deafferentation of interoceptive signals on arousal, exteroceptive perception, and (metacognitive) confidence placed in perceptual categorization. Indeed, we were able to go beyond simulated deafferentation studies and ask what it would be like if we were able to selectively lesion the precision of (i.e. confidence ascribed to) different sorts of beliefs; for example, beliefs about ‘what I am doing’, beliefs about ‘the state of the world’, and beliefs about ‘the sorts of interoceptive and exteroceptive signals I expect to encounter’.

Second, we constructed our synthetic subject in such a way that the same paradigm could be replicated in real subjects. The motivation for this is that the active inference scheme used below has an associated process theory (Friston et al., 2017a). In other words, neuronal and behavioral responses associated with inferential processes can be simulated on a trial by trial basis. This means that we can use electrophysiological, eye tracking, pupillometry and other physiological proxies to test various hypotheses that can be instantiated in the model. Crucially, this provides a link between neuronal and behavioural responses – as characterised by the latency between stimuli onset and autonomic responses (e.g., heart rate acceleration or variability) or confidence judgements (i.e., responses to how confident were you in your perceptual judgement?). In this paper, we will focus on the basic phenomenology and (some counterintuitive) results. In subsequent work, we will use this formalism to model real responses under various experimental manipulations.

In what follows, we briefly describe the generative model and inversion scheme used to simulate cardiac arousal responses. We then demonstrate the results of anatomical (deafferentation) lesions on perceptual and metacognitive behaviour, as well as simulated belief updating. Finally, we will examine the effects on synthetic heart-rate variability when changing the precision of various prior beliefs that underlie perceptual inference. We conclude with a discussion of the implications for existing research in this area – and how this research could be informed by a formal approach providing guidelines to discovery.

## Methods

### Markov Decision Process

The simulations reported below build upon the notion of active inference. This is a ‘first-principles’ approach to understanding (Bayes) optimal behaviour. Simply put, active inference treats the brain as using an internal (generative) model of the world to explain exteroceptive, proprioceptive, and interoceptive sensory data. By optimizing beliefs about variables in this model (perceptual inference), or by changing their internal or external environment (action), creatures can ensure their sensations and prior beliefs are consistent^1^. A Markov decision process (MDP) is a form of probabilistic generative model that describes the sequential dynamics of unobserved (hidden) variables (e.g., the current state of the cardiac cycle) and the sensations they cause (e.g., baroreceptor signals). The hidden variables of an MDP are hidden states (*s_τ_*) and sequences of actions or policies (*π*). The generative model then embodies the conditional dependencies between these variables, as expressed graphically in Figure 1. While we provide a brief overview here, we refer readers to (Friston et al., 2017a) for more technical detail.

**Figure 1.**
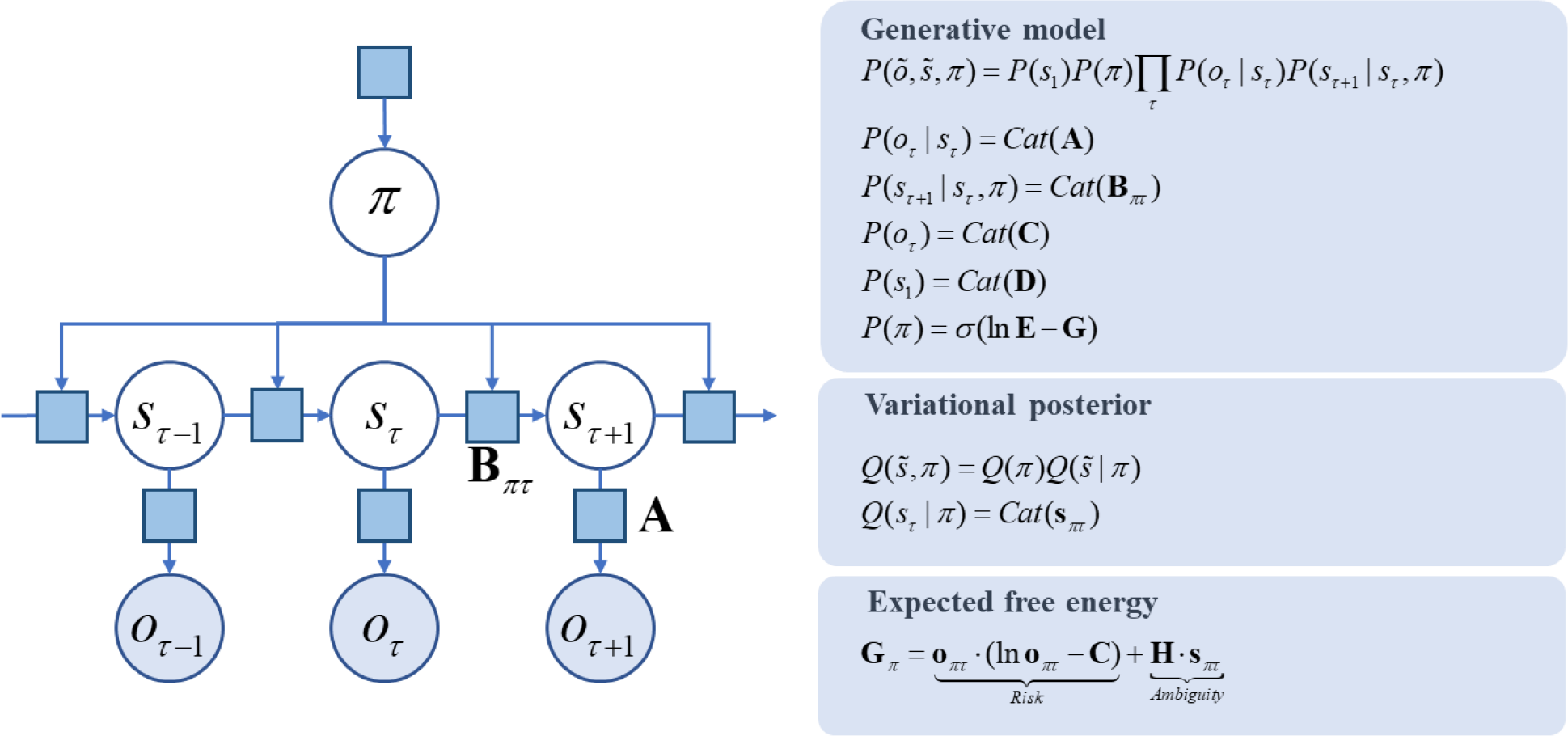
A Markov decision process generative model: the factor graph *on the left* illustrates the conditional dependencies, and independencies, between the variables in the generative model (see the main text for a description of the variables). The variables are shown in circles (with filled circles showing observable variables). An arrow from one variable to another indicates that the latter depends upon the former. The square nodes each represent probability distributions. The panels *on the right* give the forms of the distributions (associated with each square node) in the generative model, in addition to defining the expected free energy, and specifying the factorization of the approximate posterior (variational) distributions the agent possesses.

Hidden states generate observable sensory data with probabilities expressed in a likelihood matrix **A**. The states evolve through time according to a transition probability matrix, **B** and depend only on the state at the previous time, and on the policy, *π*,. Finally, we equip the generative model with preferences (**C**), prior beliefs about initial states (**D**), and prior beliefs about policies. Beliefs about policies have two parts. The first of these is a fixed bias (**E**). This may be thought of as a habit; i.e., ‘what I expect to do’ *a priori*. The second is a belief that the most probable policies are those that have the lowest expected free energy (**G**); i.e., ‘what I expect to do’ after considering the consequences of action. A simple intuition for the latter is to think of the selection between alternative courses of action as we might think of Bayesian hypothesis testing (i.e. model comparison); namely, planning as inference (Attias, 2003; Botvinick and Toussaint, 2012). Here, each policy can be thought of as an alternative hypothesis about ‘how I am going to behave’. These are evaluated in terms of prior beliefs (**E**), and the (predicted) evidence future data affords (**G**). Just as free energy is used to approximate the evidence data affords a hypothesis, expected free energy evaluates the expected evidence, under beliefs about how data are actively generated. As expressed in Figure 1, expected free energy can be separated into two parts. ‘Risk’ quantifies how far predicted observations deviate from preferred outcomes. Minimizing this ensures maintenance of homeostasis. ‘Ambiguity’ quantifies the uncertainty in the mapping from states to outcomes. Minimizing this component ensures that salient, uncertainty-resolving data are sought (leading to epistemic, information gathering, behavior).

### Synthetic Cardiac Arousal

Using the MDP scheme detailed above, we set out to simulate a cardiac arousal response to threatening stimuli (e.g., a vicious looking spider), in comparison to non-arousing stimuli (e.g., some flowers). To do this, we had to define ‘arousal’ and its interoceptive correlates. To keep things as simple as possible, we assumed the subject’s generative model included two sorts of hidden states (*interoceptive* and *exteroceptive* – and that she could adopt two modes of engagement with the world (*relaxed* and *aroused*). These sorts of generative models are generally cast as Markov decision processes, whereby transitions among (hidden) states generate observable outcomes in one or more modalities. The modalities considered here were *exteroceptive* (*non-arousing* versus *arousing* visual stimuli) and interoceptive (the cardiac phase; *diastolic* or *systolic*). Having defined the nature of the state space generating outcomes, this model can then be parameterised in a relatively straightforward fashion as outlined above. For any set of **A,B,C,D**, and **E** parameters, one can then simulate active inference using standard marginal message passing schemes (Parr et al., 2019) to optimize expectations about hidden states of the world – and the action or policy currently being pursued (technically, a policy is a sequence of actions. In what follows, we only consider policies with one action) (Friston et al., 2017a, 2017c).

Crucially, inference about policies rest upon prior beliefs that the policies will minimise expected free energy in the future. This expected free energy has both epistemic and instrumental terms; namely; the ability of any particular course of action to resolve uncertainty about hidden states (known as salience, Bayesian surprise, information gain, *etc*.) (Barto et al., 2013; Itti and Baldi, 2009; Oudeyer and Kaplan, 2009; Schmidhuber, 2010) and the pragmatic affordance (known as expected value, utility, reward, *etc.*) as specified by the prior preferences (Friston et al., 2015).

To capture the fundaments of multimodal integration – of interoceptive and exteroceptive modalities – we assumed the following, reasonably plausible, form for the model. The synthetic subject had to infer which of two policies she was pursuing: a *relaxed* policy or an *aroused* policy. These are defined operationally in terms of transitions among interoceptive states. Here, we model this in terms of two distinct forms of cardiac cycling among *diastolic* and *systolic* bodily states. When *relaxed*, the probability transitions among cardiac states meant that there were two phases of *diastole* and one of *systole*. Conversely, when *aroused*, the first *diastolic* state jumped immediately to *systole*. In brief, this means that being aroused causes cardiac acceleration and the average amount of time spent in systole. The outcomes are generated by these states were isomorphic; in other words, there was a simple likelihood mapping from states to sensations; such that the subject received a precise or imprecise interoceptive cue about the current cardiac status (i.e., *diastole* or *systole*).

On the exteroceptive side, we just considered two states of the visual world; namely, the subject was confronting an *arousing* or *non-arousing* visual object. The corresponding visual modality again had two levels (*arousing* versus *non-arousing* picture). Crucially, the fidelity or precision of this mapping depended upon the interoceptive state. When the subject was in *systole*, this mapping became very imprecise. In other words, all outcomes were equally plausible under each hidden state of the visual world. Conversely, during *diastole*, there was a relatively precise likelihood mapping. This is the crucial part of our model that links the state of the body to the way that it samples the world. Put simply, precise visual information is only available during certain parts of the cardiac cycle, which itself depends upon the state of arousal (i.e., the policy currently inferred and selected). This can be thought of as a simple approximation of cardiac and other bodily timing effects, expressed as a momentary occlusion or attenuation of sensory input by (for example) afferent inhibitory baroreceptor effects (Bonvallet and Bloch, 1961; Lacey and Lacey, 1978), or by the brief flooding of the retina during cardiac contraction.

This simple structure produced some remarkable results that speak to the intimate relationship between interoception and exteroception. These phenomena (see below) rest upon the final set of beliefs; namely, preferred outcomes. Here, the subject believed that she would be, on average, in a *systolic* state when confronted with an arousing picture and in a *diastolic* state otherwise. These minimal prior preferences then present the subject with an interesting problem. She has to choose between extending periods of precise evidence accumulation (i.e., a *relaxed* state with more *diastolic* episodes) and sacrificing precise information, via cardio-acceleration, should she infer there is something arousing ‘out there’. However, to infer what is ‘out there’, she has to resolve her uncertainty, through epistemic foraging; i.e., maintaining a *relaxed* state. We therefore hypothesised that at the beginning of each trial or exposure to a picture^2^, subjects would be preferentially in a relaxed state until they had accumulated sufficient evidence to confidently infer the visual object was *arousing* or not. If *arousing*, she would then infer herself to be aroused and enter into a period of cardio-acceleration (illustrated in Figure 3).

By carefully adjusting the precision of sensory evidence (through adjusting the **A** matrix), we could trade-off the evidence accumulation against these imperatives to simulate the elaboration of an arousing response to, and only to, *arousing* stimuli. Furthermore, we anticipated that a failure to implement a selected policy of arousal would both confound inference about the policy being pursued (i.e., an aroused state of mind) and – importantly – confidence about the exteroceptive state of affairs. The latter can be measured quantitatively in terms of the entropy or average uncertainty over hidden exteroceptive states (after taking a Bayesian model average over policies). This leads to the prediction that confidence in perceptual categorisation would not only evolve over time but would depend upon interoceptive inference. We tested this hypothesis *in silico* through various lesion experiments reported in the subsequent sections (Figures 3 - 5). In what follows, we illustrate the belief updating and arousal responses under ‘normal’ priors (i.e. precisions) based upon the generative model above (summarized graphically in Figure 2).

**Figure 2:**
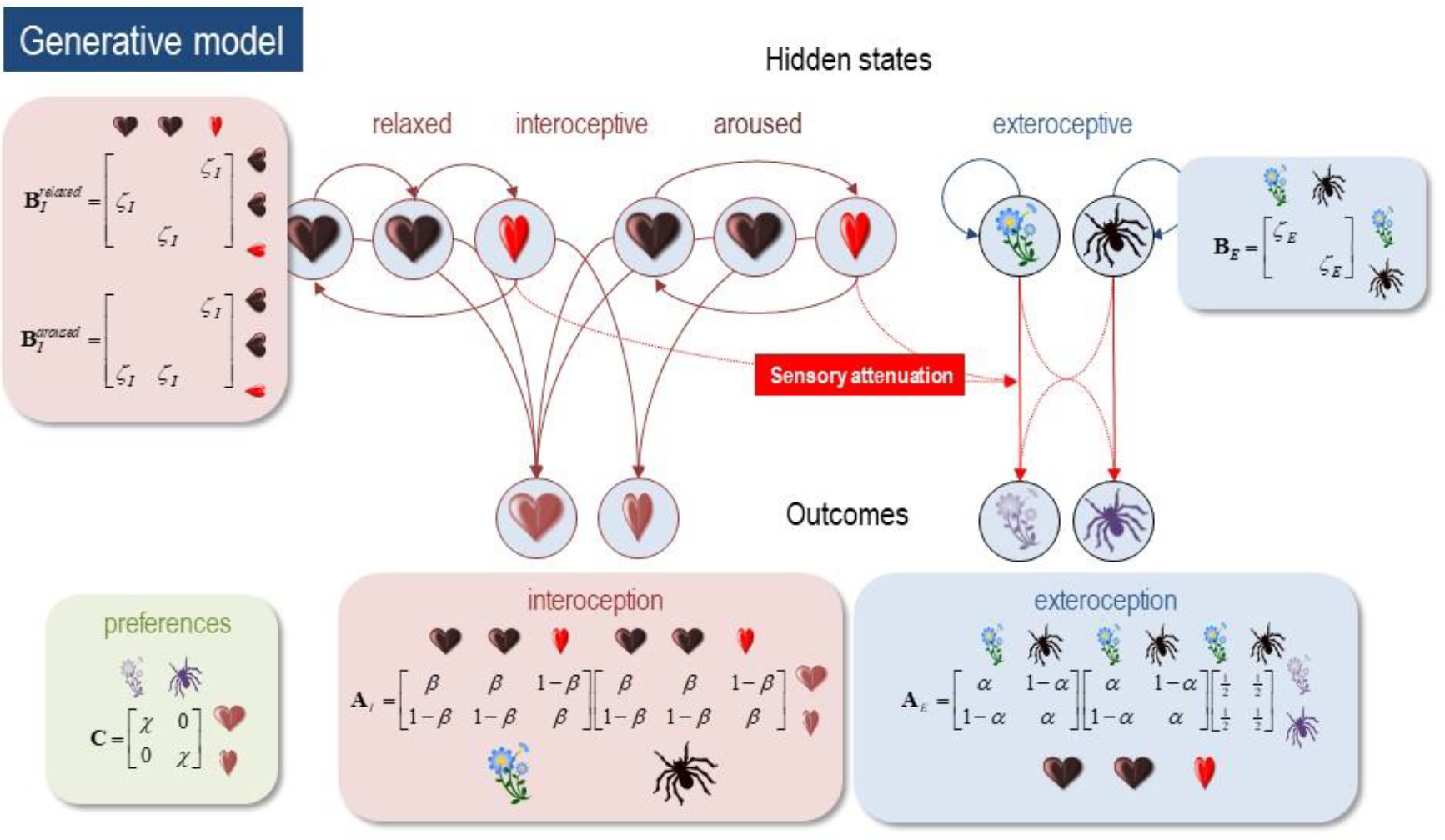
*the generative model*. This schematic illustrates how hidden states cause each other and sensory outcomes in the interoceptive and exteroceptive domain. The upper row describes the probability transitions among hidden states, while the lower row specifies the outcomes that would be generated by combinations of hidden states that are inferred on the basis of outcomes. The green panel specifies the models prior preferences; namely, the sorts of outcomes it expects to encounter. Please see main text for a full explanation. Although this figure portrays interoceptive and exteroceptive outcomes as separate modalities, they were in fact modelled as combinations – so that the prior preferences could be evaluated (this is necessary because the preferred physiological outcome depends upon the visual cue). In this model, the precisions are denoted by Greek letters and control the fidelity of various probabilistic mapping is (i.e., the likelihood or **A** matrices and the transition or **B** matrices).

**Figure 3.**
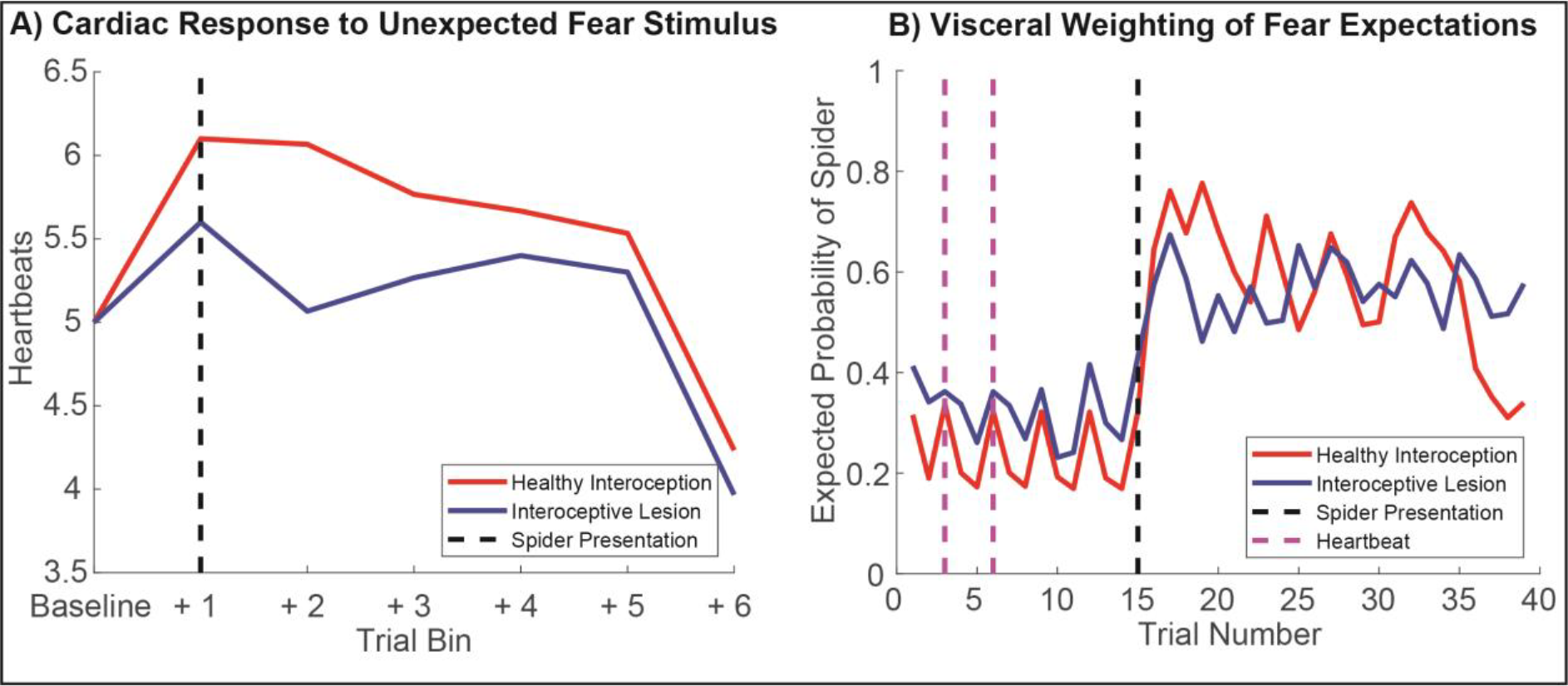
Simulated Physiology and Perceptual Inference. To establish the face validity of our model, we first set out to reproduce some basic psychophysiological phenomenology and establish how these phenomena change under ‘healthy’ (i.e., normative) versus ‘visceral lesion’ parameter settings. To do so, we fed agents a fixed sequence of cardiac and exteroceptive stimuli, such that the first 14 trials constituted a ‘baseline’ period of cardiac quescience (i.e., a steady heart rate), in the absence of arousing stimuli. On the 15th trial, an unexpected arousing stimulus (a ‘spider’) is presented and a further 85 trials simulated. This simulation was repeated for 60 simulated participants, each with randomized starting values, half of which had ‘lesioned’ interoceptive precision (β = 0.5, blue lines). Under these conditions our synthetic subjects exhibit a clear ‘startle’ or ‘defense’ reflex (Graham and Clifton, 1966; Sokolov, 1963), characterized by an immediate cardio-acceleration (left panel) and a dramatic shift in the posterior expectation of encountering another threatening stimulus. Interestingly, during the baseline period the posterior expectation of encountering a threat stimulus oscillates with the heartbeat; i.e., the lesioned subjects show both an attenuation of the cardiac response and a blunted belief update. Note that for the right panel, only trials 1-40 are shown. On the left, blue lines show summed heartbeats (time spent in systole) for 15-trial bins; on the right, lines depict the median posterior probability that the agent will see a spider on the next trial. See *Methods* and *Results* for more details.

### Simulations

We implemented a minimal model of interoceptive and emotional inference – in the sense that one’s state of active engagement with the world may be inferred from its interoceptive and exteroceptive consequences. In this minimal model, the two domains of perception are coupled by – and only by – sensory attenuation: i.e., attenuation of sensory precision in the visual domain during (inferred) systole. Precision refers to the reliability or confidence ascribed to a given probabilistic belief. Within this model there are four kinds of precision; namely, sensory precision in the visual (α) and cardiac (β) domains and the precision of state transitions among interoceptive and exteroceptive states. For clarity, we will refer to the precision of transitions as (inverse) *volatility* and use *precision* to refer to sensory (i.e. likelihood) precision. In this example, because there are only two states, the corresponding parameters of the generative model control both the expected contingencies and their precision. In other words, when α (or β) decreases to 1/2, sensory signals become imprecise and completely ambiguous. In what follows, we will focus on manipulations of precision under a canonical volatility of ζ = 0.9. In other words, we will assume that our synthetic subject believes state transitions among phases of the cardiac cycle follow each other fairly reliably with a 90% probability. Similarly, if there is a flower ‘out there’, then there is a 90% probability that it will remain there at the next sample. Cardiac and visual stimuli were generated by the same precisions and volatilities as assumed by the subject’s generative model.

We conducted three sets of simulations to illustrate the sorts of behaviours that emerge under this active inference scheme – and to establish the construct validity of the model in relation to empirical phenomena that speak to the influence of interoception on exteroception and *vice versa*. This enabled us to illustrate the basic phenomenology of our agent – in terms of simulated perceptual inference and cardiac physiology – under some differing levels of sensory precision.

In the first set of simulations (Fig. 3), we focused on the physiological and psychological response to arousing stimuli. To do so, we tested the hypothesis that the unexpected presentation of a ‘spider’ would induce an aroused state – as reflected in an increased heart rate – and a greater posterior expectation of encountering an arousing spider stimulus on the next trial. To evaluate this hypothesis, we supplied the subject with a fixed sequence of 15 stimuli – in both the cardiac and visual domains – and examined the posterior beliefs about the next exteroceptive state following a period of relaxed cardiac input. Note that this is possible precisely because our generative model includes beliefs about the future – including the next hidden state and subsequent sensory sample. Here, we used as outcome measures the agent’s evoked cardiac acceleration response (calculated by binning the number of siastole events across the experiment) and the agent’s posterior belief that the next stimulus would be threatening. These simulations were repeated 60 times with randomized starting values, such that the first thirty ‘healthy’ agents where compared to an ‘interoceptive lesion’ group for whom interoceptive precision had been attenuated (β = 0.5). This enabled us to not only establish the interaction of fear expectations and cardiac arousal, but also to demonstrate how these responses change when interoceptive sensory precision is ablated.

In the second set of simulations (Fig. 4), our focus moved from perceptual to metacognitive inference. Here, we examined the interaction between exteroceptive and interoceptive sensory precision on the one hand and their coupling to cardiac timing and metacognition (posterior confidence) on the other. Our goal here was to illustrate how both interoceptive and exteroceptive precision interact to influence metacognitive inference, and to link these to empirical findings showing that cardiac arousal biases metacognition (Allen et al., 2016b; Hauser et al., 2017a). For these, we used the uncertainty about inferred exteroceptive and interoceptive states (as quantified by the summed entropy of posterior beliefs for each state) as outcome measures, simulated under a range of cardiac and visual precision settings (figure 3A). To further illustrate how these effects oscillate with the cardiac rhythm, we separated these measures for each phase of the cardiac cycle (early diastole, late diastole, systole). We then repeated these analyses comparing ‘healthy’ interoceptive inference agents (α & β = 0.9), to agents for whom either exteroceptive or interoceptive precision was lesioned (α or β = 0.5, respectively). In virtue of our coupling of exteroceptive sensory precision to the cardiac cycle, we anticipated that metacognitive confidence (outscored by the negative entropy of posterior beliefs) would depend on the precision of both interoceptive and exteroceptive states, and that this effect would clearly oscillate with the cardiac cycle. Further, we expected in the extreme case of our ‘lesioned’ subjects, these effects would be further exacerbated such that interoceptive and exteroceptive uncertainty would increase dramatically, under their respective lesion conditions.

**Figure 4.**
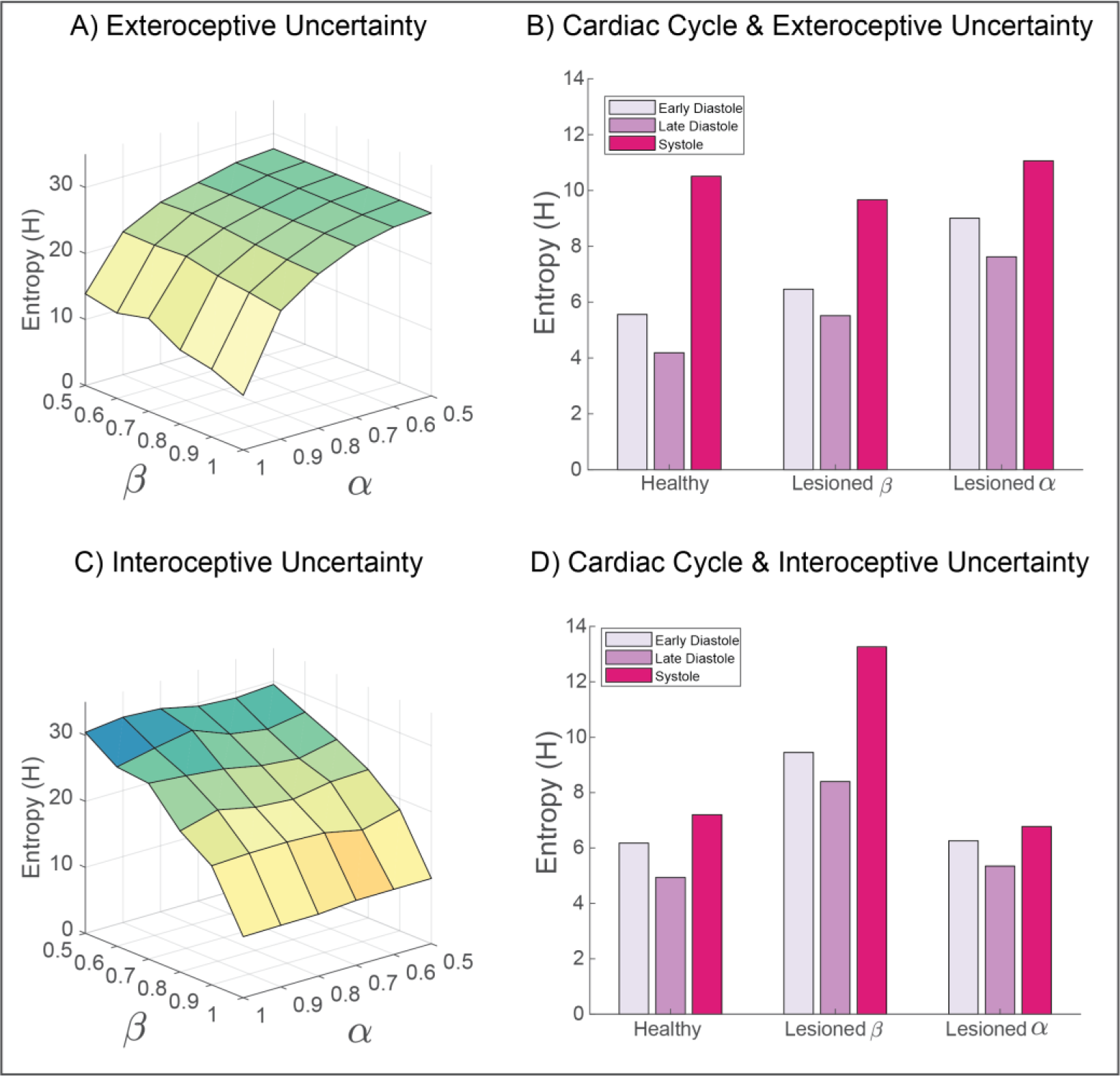
Simulating the influence of interoceptive and exteroceptive precision on metacognitive uncertainty. To explore how interoceptive inference influences metacognition, we measured the summed entropy of beliefs for both exteroceptive (top panels) and interoceptive (bottom panels) states. By simulating the full range of sensory precision values, from lesioned precision (α or β = 0.5) to ‘hyper-precision’ (α or β = 1), the predominant pattern of interactions is revealed. **A)** For exteroceptive inferences (i.e., the agent’s belief that a spider or flower is present), the principle entropy gradient is characterized by reductions in exteroceptive precision. This effect is modulated in part by interoceptive precision; for example, the lowest uncertainty is obtained when interoceptive and exteroceptive precision are maximal. **B)** Separating exteroceptive uncertainty by each phase of the cardiac cycle reveals a clear effect of the heartbeat on belief entropy, which is modulated most strongly by lesioning the precision of exteroceptive predictions. Lesioning interoceptive uncertainty does raise the overall level of exteroceptive uncertainty, but to a lesser degree. Note that altering exteroceptive precision only affects the diastolic phases (as precision is already attenuated during systole). Interoceptive lesions preclude precise inferences about the cardiac phase, so reduce the discrepancy in uncertainty between these phases. **C)** Similar to exteroceptive belief, interoceptive metacognition is predominately influenced by interoceptive precision. **D)** The cardiac cycle also modulates the overall uncertainty of interoceptive beliefs; this effect is greatly increased when interoceptive precision is lesioned. Interestingly, exteroceptive lesions primarily reduce the differentiation between cardiac states.

Finally, to complement these simulations we modelled the response of first and second order statistics of the physiological responses to changes in sensory precision. These were based upon simulated heart rate (frequency of systole) and the heart rate variability (HRV) assessed over multiple trials or heartbeats (Fig. 5). Our objectives here were; 1) to test the hypothesis that fluctuations in both low-and high-frequency synthetic heart rate variability can be produced by altering the balance of interoceptive sensory precision versus the prior precision for the aroused sympathetic policy, and 2) to illustrate how generative modelling of interoceptive active inference can be used to phenotype maladaptive inference parameters from observed heart-rate data (i.e., interoceptive inference phenotyping). For this analysis, we simulated 1000 trials under three canonical parameter settings designed to resemble potential neuropsychiatric phenotypes of interest: healthy interoception (α = 0.8, β = 0.8, prior probability of parasympathetic policy = 55%), hyper-precise interoceptive sensation (α = 0.8, β = 1, prior probability of parasympathetic policy = 55%), and hyper-precise arousal priors (α = 0.8, β = 0.8, prior probability of sympathetic policy = 75%).

**Figure 5.**
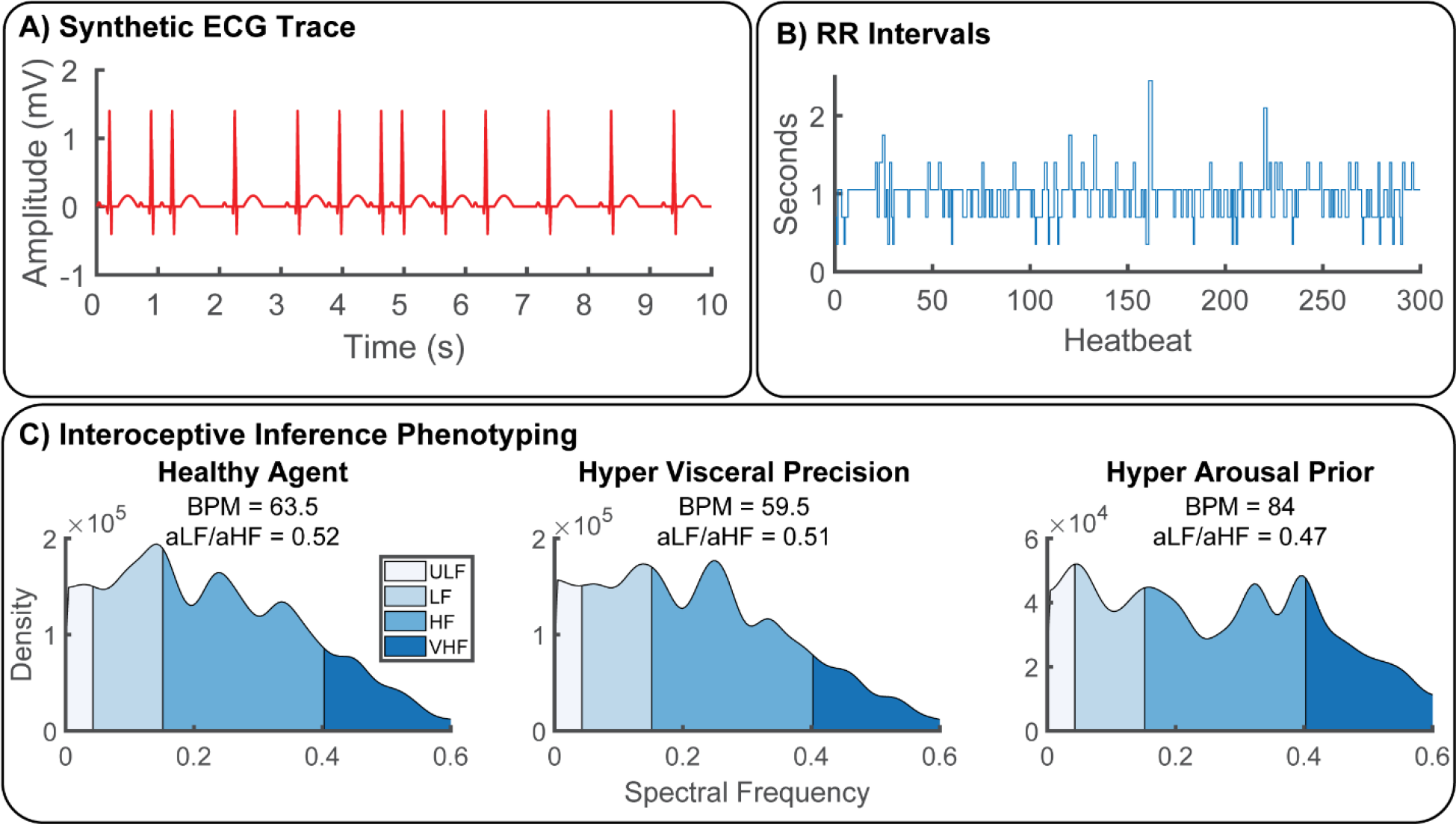
Synthetic Heart-Rate Variability (HRV) and Interoceptive Computational Phenotyping. To illustrate the potential of our approach as a generative model of physiological reactivity, we produced synthetic heartbeat traces and analyzed these with a standard time-frequency approach under various canonical parameter settings. **A)** Synthetic ECG traces produced by convolving a standard QRS-wave function with systole events generated by our model. **B)** These where then transformed into RR-intervals by assuming an 350ms sampling rate, **C)** Power spectra of RR-intervals were calculated using Welch’s method and categorized as ultra-low (ULF), low, (LF), high (HF), and very high frequency (VHF) bands for each simulated agent. Physiological responses were then summarized in terms of beats-per-minute (BMP) and sympathovagal balance (ratio of area under curve for each frequency band, aLF/aHF) (Malliani et al., 1991). To illustrate the potential of our approach for interoceptive computational phenotyping, we simulated three different agents – one with healthy interoceptive inference (bottom left), another with hyper-precise visceral sensations (bottom middle), and another with hyper-precise priors for the aroused (sympathetic) policy (bottom right). These each produce unique interoceptive inference ‘fingerprints’; i.e., the individual patterns of heart-rate variability produced by these parameter settings. In this example, hyper-precise visceral sensations reduce heart-rate and shift overall peak frequency to the high-frequency domain, whereas hyper strong arousal priors induce strong heart-rate acceleration coupled with attenuated ultra-low and ultra-fast oscillations. In the future, these idiosyncratic patterns could be used to identify maladaptive interoceptive inference from heart-rate data.

The resulting time-series of systole events from each agent were then convolved with a canonical QRS-wave response function and transformed into normalized beat-to-beat RR-intervals. To normalize the (arbitrary) sampling rate of each time-series, we assigned a 350ms repetition time (TR) for each state of the MDP simulation, such that the healthy agent had a heart rate of approximately 60 BPM. The time intervals between successive synthetic R-peaks was then calculated. As the RR interval data is unevenly sampled, the time series was linearly interpolated. The power spectrum was then estimated using Welch’s method. In line with conventional HRV analysis, the power spectra were then categorized into four frequency bands corresponding to ultra-low (0 - 0.04 Hz), low (0.04- 0.15 Hz), high (0.15 - 0.4 Hz), and very-high (> 0.4 Hz) frequency categories. Finally, to summarize the physiological response of each agent, we calculated the beats per minute (BPM) and the ratio between low and high frequency components (LF/HF), i.e., sympathovagal gain or balance. Sympathovagal valance is thought to index the balance of sympathetic and vagal outflows and is frequently interpreted implicated in stress and other psychophysiological and clinical disorders (Malliani et al., 1991; Strigo and Craig, 2016) but see (Eckberg Dwain L., 1997; Heathers, 2012) for critique. Sympathovagal balance was calculated as the ratio of area under the curve (AUC) for low and high-frequency HRV; AUC^LF^/ AUC^HF^.

## Results

### Simulated Physiology and Perceptual Active Inference

To establish the face validity of our model, we simulated the basic psychophysiological behavior of our active inference agent. This involved simulating a fixed series of stimuli (states) in which the heartbeat was forced to remain relaxed – and only non-arousing (flower) stimuli were presented. On the 15th trial, an unexpected spider stimulus was presented, and the simulation continued for a further 85 trials. Thus, by evaluating the evolution of the agent’s synthetic interoceptive physiology and exteroceptive beliefs, before and after the quiescent baseline period, we hoped to reproduce and illuminate well-known psychophysiological phenomenon such as the defensive startle reflex (Graham and Clifton, 1966; Sokolov, 1963).

This analysis, illustrated in Figure 3, revealed several interesting aspects of interoceptive active inference. Over 60 simulations there was a clear and robust increase in heart-rate acceleration, following the presentation of the unexpected or novel threat stimulus. During subsequent experiences of its own heartbeat and spiders or flowers, this response habituates, resulting in a gradual heart-rate deceleration from the evoked cardiac response. This robust modulation of heart-rate was accompanied by a jump from an expected probability of encountering a spider of about 25% to almost 65% following the spider presentation. This combined response of both the heartbeat and fear-expectations is further underscored by the curious oscillation of cardiac states and the expected probability of observing a spider; note the uptick in expectations of approximately 5% on each systole event (denoted by the pink dotted line on Figure 3, right panel). A simple explanation for this result is that, during presentation of a stream of flowers, we can confidently infer a safe external environment. This accounts for the relatively low probability of spiders in the earlier part of the plot. However, during systole, attenuated integration of exteroceptive data leads to greater uncertainty. Going from a confident inference in the absence of a spider to a more uncertain inference necessarily increases the probability of a scary environment during this cardiac phase. This offers a simple perspective on previous experimental work suggesting that fear-stimuli are potentiated when presented in synchrony with the heart (Garfinkel et al., 2014; Garfinkel and Critchley, 2016); namely, that a mechanism underlying this effect can be found in the link between cardiac active inference and fear expectations. In short, under generative models of an embodied world – in which sensory sampling depends upon fast fluctuations in bodily states – there is a necessary dependency of Bayesian belief updating (i.e., perceptual inference) across all modalities on introception.

When comparing these effects in the healthy agent to our sample of ‘lesion patients’, a few sensible but counter-intuitive consequences ensue. In the physiological domain, when presented with the unexpected arousal stimulus, the lesioned agent shows a blunted cardiac acceleration response, which remains diminished throughout the simulated trials. This blunting effect is mirrored for fear expectations in the immediate post-stimulus (e.g., trials 15-20) period, further underlining the close link between visceral and exteroceptive inference in our agent. The reason for this blunting likely results from the differing exteroceptive precision anticipated during different cardiac phases (see also Fig. 4B). A visual impression – consistent with a spider – is highly informative during diastole but must be treated with suspicion during the sensory-attenuated systolic phase. This implies a blunting of belief-updating in response to a spider, when we are unsure of cardiac phase (compared to when we are confident of a diastolic phase). A further interesting result is found when examining the controlled baseline period (trials 0-15); baseline fear expectations in the interoceptive lesion group are actually slightly enhanced by about 5-10% posterior probability. This lends an interesting embodied twist to the literature on ‘circular inference’, psychosis and hallucinations (Denève and Jardri, 2016; Powers et al., 2017), suggesting that the disruption of interoceptive precision may be one mechanism underlying hallucinations, particularly those that are affective and/or somatic in nature.

### Simulating the Influence of Sensory Precision on Metacognition

We next performed a series of simulations to tease apart how interoceptive and exteroceptive precision (and their disruption) influence ‘metacognition’; that is the uncertainty in our agent’s beliefs. To do so, we first measured the Shannon entropy for interoceptive and exteroceptive inferences (summed across both factors of posterior beliefs) under a full range of precision settings from 0.5 - 1. To highlight the oscillatory nature of cardiac effects, we then calculated the same entropy measure separately for each cardiac state (early diastole, late diastole, systole). Finally, we compared these ‘healthy’ simulations to extreme degradations in sensory precision (exteroceptive and interoceptive ‘lesions’), to better understand how disruptions of each modality are integrated in metacognition.

This analysis revealed first of all that, in our simplified model, metacognitive uncertainty is largely influenced by the unimodal precision of each domain. For both exteroceptive and interoceptive inferences, the slope of the uncertainty gradient (Fig. 4A & C) was predominantly characterized by degradations in the precision of the corresponding modality. However, this modularity is not complete; exteroceptive uncertainty is at its lowest when interoceptive and exteroceptive precision are maximal. Similarly, although interoceptive uncertainty is largely driven by interoceptive precision, small interactions with exteroceptive precision can be observed in the plotted uncertainty gradient. One interesting isomorphism, however, is that overall interoceptive uncertainty is less affected by exteroceptive precision. This is likely due to that fact that in our model, the cardiac cycle directly modulates exteroceptive precision, whereas exteroceptive states only indirectly modulate interoceptive responses, via policy selection.

This intricate relationship of the cardiac cycle and metacognitive uncertainty is further teased apart in Figure 4B, which shows clearly that exteroceptive confidence oscillates with each phase of the heartbeat, being highest at diastole. This is an unsurprising feature of our model: on each diastole, phase exteroceptive sensory precision drops effectively to null. Interestingly however, average exteroceptive uncertainty is modulated in a fairly linear fashion by visceral and exteroceptive lesions: average entropy is increased modestly by lesioning interoceptive precision and more robustly by exteroceptive lesions. Whereas interoceptive lesions caused the greatest increase in interoceptive entropy, exteroceptive lesions seem to exert a specific effect of unbinding entropy from the individual cardiac state, again mirroring the isomorphic representation of these states in uncertainty. This is a sensible finding, as the manipulation leads to relatively high uncertainty in the mapping between hidden states and outcomes during all cardiac phases, not just during the previously attenuated systolic phase. This sort of chronic hypo-arousal – as a consequence of a failure to contextually modulate precision – is not unlike that which may underwrite the negative symptoms of schizophrenia or depression.

### Synthetic Heart-Rate Variability (HRV) and Embodied Computational Phenotyping

In our final set of simulations, we illustrated how the interoceptive inference approach developed here offers a new means for analyzing and interpreting fluctuations in observed physiological data. Our goal here was to demonstrate the potential for generative modelling and ‘embodied computational phenotyping’; i.e., the identification of specific parameters of brain-body interaction underlying maladaptive interoceptive inference in psychiatric and other health-harming disorders; e.g., (Peters et al., 2017).

To this end, we generated synthetic cardiac data by convolving our train of cardiac events with an ECG response waveform. Following standard methods, we then calculated the normalized beat-to-beat intervals and performed a time-frequency analysis of the resulting RR-interval data. By repeating this analysis for a ‘healthy’ agent under normative values, an agent with interoceptive ‘hyper-precision’ (i.e., β = 1), and an agent with an overly precise prior beliefs about its own arousal, we illustrate how individual HRV fingerprints are linked to unique patterns of interoceptive active inference.

This analysis showed that, despite the exceedingly simple (biomechanically speaking) conditions of our model, sensible and interesting patterns of heart-rate variability emerge for different combinations of interoceptive sensory and prior precision. Specifically, we found that whereas the healthy agent exhibited a relatively relaxed profile in terms of heart rate and sympathovagal balance (BMP = 63.5, peak frequency = 0.14 Hz) – predominated by low versus high frequency oscillations (aLF/aHF = 0.52) – an agent with hyper-precise visceral sensations exhibited a mild downshift in heart-rate coupled (BPM= 59.5) with an overall increase in high-frequency oscillations (aLF/aHF = 0.51, peak frequency = 0.25). In contrast, the agent with hyper-precise arousal priors showed a strong bimodal modulation of both ultra-low and ultra-high frequencies HRV (peak frequencies = 0.04 Hz & 0.34 Hz, respectively), coupled with a strong increase in heart-rate (BPM = 84) and high versus low-frequency outflow (aLF/aHF = 0.47). These results speak to the unique role of different active inference parameters in producing highly idiosyncratic patterns of HRV variability. In the future, our model may be enhanced to subserve computational phenotyping of individual differences and/or patient subgroups categorized by the balance of visceral precision and arousal policy priors from raw HRV data alone.

## Discussion and Conclusions

In the present work, we have introduced the first formal model of interoceptive inference as applied to emotion, exteroceptive perception, and metacognitive uncertainty. Through a variety of simulations, we demonstrated that this model can reproduce a variety of psychological and physiological phenomena, each of which speak to a unique domain of the burgeoning interoceptive inference literature (Allen and Friston, 2018; Feldman and Friston, 2010; Seth, 2013), and the application of interoceptive inference to computational psychiatry (Owens et al., 2018; Petzschner et al., 2017). This formulation of interoceptive inference reproduces some of the finer details of physiological responses to arousing stimuli that, crucially, are emergent properties under the simple assumption that people use generative models to infer the state of their lived world.

The form of the generative model and (neurobiological implausible) belief updating used in this paper are generic: exactly the same scheme has been used to simulate a whole range of processes, from neuroeconomic games to scene construction and attentional neglect (Friston et al., 2017a; Parr and Friston, 2018). The key aspect of the generative model introduced here is that the quality (i.e., precision) of sensory information depends upon fluctuations in (inferred) autonomic states. This simple fact underwrites all of the phenomenology illustrated above; both in terms of simulated physiology and accompanying belief updates. The explicit inclusion of interoception into active inference licenses us to talk about ‘fear’ and in the sense that affective inference is thought to emerge under models that generate multimodal predictions that encompass the interoceptive domain. Furthermore, casting everything as inference enables a metacognitive stance on belief updating, in the sense that one can quantify uncertainty invested in beliefs about states of the body, states of the world and, indeed, states of (autonomic) action.

In particular, we show that by simulating periodic attenuation of exteroceptive sensory inputs by the cardiac cycle, affective expectations become intrinsically linked to afferent interoceptive signals through a startle reflex-like phenomenon. This linkage not only induces oscillatory synchrony between the heartbeat and exteroceptive behavior, but also propagates to metacognitive uncertainty (i.e., the entropy of posterior beliefs). This latter finding speaks to numerous reports of metacognitive bias (e.g., confidence-accuracy dissociation) by illustrating how the precision of interoceptive states can directly influence exteroceptive uncertainty (Allen et al., 2016b; Boldt et al., 2017; Spence et al., 2016). By simulating synthetic heart-rate variability (HRV) responses, we further illustrated how idiosyncratic patterns of aberrant interoceptive precision-weighting can be recovered through generative modelling of physiological responses, opening the door to computational phenotyping of disordered brain-body interaction in the spirit of (Schwartenbeck P and K Friston 2016). In what follows, we outline some of what we view as the most promising future directions for this work, sketch a proposed neuroanatomy underlying our model, and point out a few limitations for consideration.

By focusing on the periodic nature of the cardiac cycle, and concomitant influences on exteroceptive perception, our goal was to provide an initial proof-of-principle, illustrating how visceral and exteroceptive signals may be combined under active inference. Our aim was not to suggest that our model provides the ultimate view of interoceptive inference; indeed, we view the present work as a starting point that can be taken forward in a variety of research directions, some of which are outline below.

In this paper, we formalized the hypothesis that frequently reported effects of cardiac timing on perception could arise as a function of periodic sensory attenuation – but the reader should feel encouraged to test their own hypotheses within the openly available MDP framework. Our intention here was also not to prioritize cardiac-brain interaction over e.g., gastric or respiratory cycles, but instead to provide a toy example, to show how these systems may be subjected to formal analyses. This was motivated by the large predominance of research on cardiac-brain interaction; however, we do anticipate that the periodic attenuation of sensory precision by visceral signals is likely to provide a general explanation of brain-body interaction.

Neurophysiologically, the principal means by which cardiac signals influence the central nervous system is through the afferent cardiac baroreceptors. These pressure-sensitive neurons, located primarily in the aorta and carotid artery, are triggered by the systolic pressure wave generated when the heart contracts. Far from being restricted to homeostatic function only, it was first reported (nearly a century ago) that afferent baroreceptor outputs induce a general inhibitory effect on cortical processing (Bonvallet et al., 1954; Bonvallet and Bloch, 1961; Koch, 1932). These findings were later extended by Lacey and Lacey (1978) who proposed the “neurovisceral afferent integration hypothesis”, positing that cardiac acceleration and deceleration serve to respectively disengage or engage with an exteroceptive stimulus via cortical inhibition.

In parallel, the soviet psychologist Evgeny Sokolov proposed that novelty (but not threat) evoked heart-rate deceleration was a core component of the ‘orienting reflex’ (Sokolov, 1963). By reducing overall cardiac output, this reflex served to limit the contribution of cardiac signals to cortical noise boosting overall signal-to-noise ratio^3^. In contrast, Sokolov theorized that the defensive startle reflex – in which an extremely strong (e.g., the loud bang of a starting gun) or unexpectedly aversive (e.g., the sudden presentation of a spider) stimulus evokes cardiac acceleration – facilitated the disengagement of cortical processing, to initiate fight-or-flight responses. These theories in turn sparked a wave of empirical studies attempting to link cardio-acceleration and deceleration responses to increased or decreased exteroceptive sensitivity, which continues to this day (Azevedo et al., 2017; Cohen et al., 1980; Delfini and Campos, 1972; Edwards et al., 2009; Elliott, 1972; Garfinkel et al., 2014; Ghione, 1996; Park et al., 2014; Salomon et al., 2016; Sandman et al., 1977; Saxon, 1970; Velden and Juris, 1975).

While these findings highlight the intricate relationship between cardiac timing and exteroceptive psychophysics, so far a consistent pattern of findings (e.g., sensory signal enhancement and/or inhibition) has failed to emerge (see Elliott, 1972 for one critique). A cursory review of this literature reveals evidence for both exteroceptive enhancement and suppression, depending upon the specific nature of the exteroceptive stimuli (i.e., whether they are inherently aversive, sociocultural, or neutral in nature), the context of the arousal (including specific stimulus and response timing), and other psychophysiological moderators; such as age, gender, and overall physical fitness. Accordingly, more recent proposals have focused on more modality-specific exteroceptive enhancement by cardiac signals. For example, that cardiac-exteroceptive effects specifically potentiate fear or threat signals (Garfinkel and Critchley, 2016) or the generation of a subjective first-person viewpoint (Park and Tallon-Baudry, 2014).

We offer a unique synthesis of these views, expressed in terms of interoceptive inference. In our model, the cyclic influence of the heart on exteroception is exerted primarily through the attenuation of sensory precision on each systolic contraction, which in turns influences the selected (multimodal) arousal policy as determined by the agent’s preferences. The coupling of sensory attenuation to the cardiac cycle endorses the notion that baroreceptors exert an inhibitory influence on the brain. Beyond this direct effect, our model can also be understood in light of the well-known relationship between intrinsic noise fluctuations in the brain and cardio-respiratory cycles (Birn, 2012; Karavaev et al., 2018). Physiological oscillations exert non-neuronal influences on spontaneous brain activity via a variety of more or less direct causal influences; for example, at each heart beat visual input to the retina is briefly attenuated by a pulsatile blood inflow. Similarly, with each cardio-respiratory cycle, fluctuations in cerebral pulsatile motion and blood pressure induce neurons to spontaneously fire, shaping the ‘infraslow’ brain dynamics (Golanov et al., 1994; Karavaev et al., 2018; Zanatta et al., 2013) that influence the overall global dynamics of neural excitability and connectivity (Fox et al., 2007, 2006; Fox and Raichle, 2007). Our suggestion is that, insofar as the brain must model its own dynamic noise trajectories as a function of active self-inference, non-neuronal sources of variability such as inscribed by visceral rhythms must be incorporated within the brain’s generative model of its own percepts. Interoceptive fluctuations are thus an important influence over the precision of exteroceptive sensory channels, and interoception is itself the means by which the brain infers and controls its own pathway through these precision trajectories. The modelling introduced here can thus be expanded beyond the cardiac domain to the more general problem of modelling how spontaneous fluctuations in neurovisceral cycles (including heart-rate variability) influence information processing and behavior.

What then, explains the lack of consistent results within the cardiac timing literature? In contrast to the binary on/off hypotheses proposed by Lacey or Sokolov, our simulations highlight the context-sensitive manner by which ascending visceral signals modulate the precision of both interoceptive and exteroceptive inferences. For example, our simulation of the startle response (illustrated in Fig. 3) clearly indicates that the functional impact of cardio-ballistic responses is coupled to the agent’s baseline prior expectations, as well as the overall precision of active inference and policy selection. In this sense, whether a specific cardiac response is likely to potentiate or inhibit a specific domain (e.g., fear) depends upon the specific weighting of arousal policy priors, the precision of incoming exteroceptive and interoceptive sensations, and the linkages thereof as determined by the task itself. In other words, the specific balance of prior beliefs and sensory information, in a given cognitive or affective domain, must be addressed before one can predict the exact directionality of an interoceptive effect on perception, or *vice versa*. Here, we modelled the generation of arousal policies as a function of hyper-parameters governing the preferred policy. In the future this can be unpacked further by examining the divergence between prior and posterior beliefs about these policies (e.g., through inferred epistemic value). Through Landauer’s principle, this divergence may be equated with the associated metabolic costs of computation and the conceptual notion of interoceptive self-modelling (Kiverstein, 2018; Limanowski and Blankenburg, 2013; Seth and Tsakiris, 2018).

### The computational neuroanatomy of interoceptive inference

Having addressed the construct validity of our model, we now speculate as to some likely neuronal substrates of the message passing implied by variational inference. Interoceptive inference can be broken down into four core functional domains: basic sensory-motor control, conscious interoceptive (perceptual) awareness, metacognitive monitoring, and hedonic (intrinsic) value. In our model, we focused primarily on the simplest possible implementation of interoceptive inference, corresponding to the sensory-motor domain (i.e., ascending and descending cardiac pathways) and their low-level interaction with exteroceptive inference, via neuromodulatory gain control. Future work will benefit from expanding upon our representation of uncertainty to include the computation of epistemic and/or intrinsic value as proxies for these higher-order interoceptive systems (Friston et al., 2017b; Parr and Friston, 2017).

Accordingly, in our sketch of the putative neuroanatomy underlying cardiac active inference (Fig. 6), we focus primarily on the neuronal substrates that inscribe low-level viscerosensory and visceromotor control, as well as some hierarchically superior regions related to emotional salience and interoceptive awareness. For simplicity, our model depicts only the minimal neuronal message passing scheme implied by our generative model; as such, we have omitted many of the intermediary relay nodes; e.g., in the thalamus and ventral visual stream. Afferent baroreceptor signals are transmitted along the ascending vagus to the rostrum of the nucleus tractus solitarus (NTS, Mifflin and Felder, 1990; Miura and Reis, 1972). From here, ascending viscerosensory signal are projected via brainstem and midbrain nuclei to the thalamus, somatosensory cortex, and posterior insula (Cechetto and Saper, 1987; Craig, 2002); ascending cardio-sensory outcomes are thus encoded in the NTS and then passed to the posterior insular cortex (PIC) as inferred interoceptive states. The PIC has a well-known role as primary viscerosensory cortex; electrical stimulation of this area elicits phantom visceral sensations (e.g., pain, heart-rate acceleration) (Chouchou et al., 2019; Oppenheimer et al., 1992) and bolus isoproterenol infusions increase the intensity of cardiorespiratory sensations and concomitant PIC activations (Hassanpour et al., 2016; Khalsa et al., 2009). In parallel, visual sensory outcomes are passed via the second cranial nerve to the superior colliculus, where they inform exteroceptive inference in the amygdala, which is well-situated to process salient emotional stimuli (Anderson and Phelps, 2001; Liddell et al., 2005). These interoceptive and exteroceptive expectations then converge in the anterior insular cortex (AIC), where they inform the selection of the appropriate autonomic policy. Finally, the selected policy is passed down the hierarchy via descending pathways (likely carried by von Economo neurons), to eventually engage the rostral nucleus ambiguus and descending vagus, decelerating the heart-rate when the relaxed policy is selected. Collectively, the scheme represents a multimodal reflex arc interlinking exteroceptive and interoceptive domains to specific patterns of cardio-ballistic responses.

**Figure 6.**
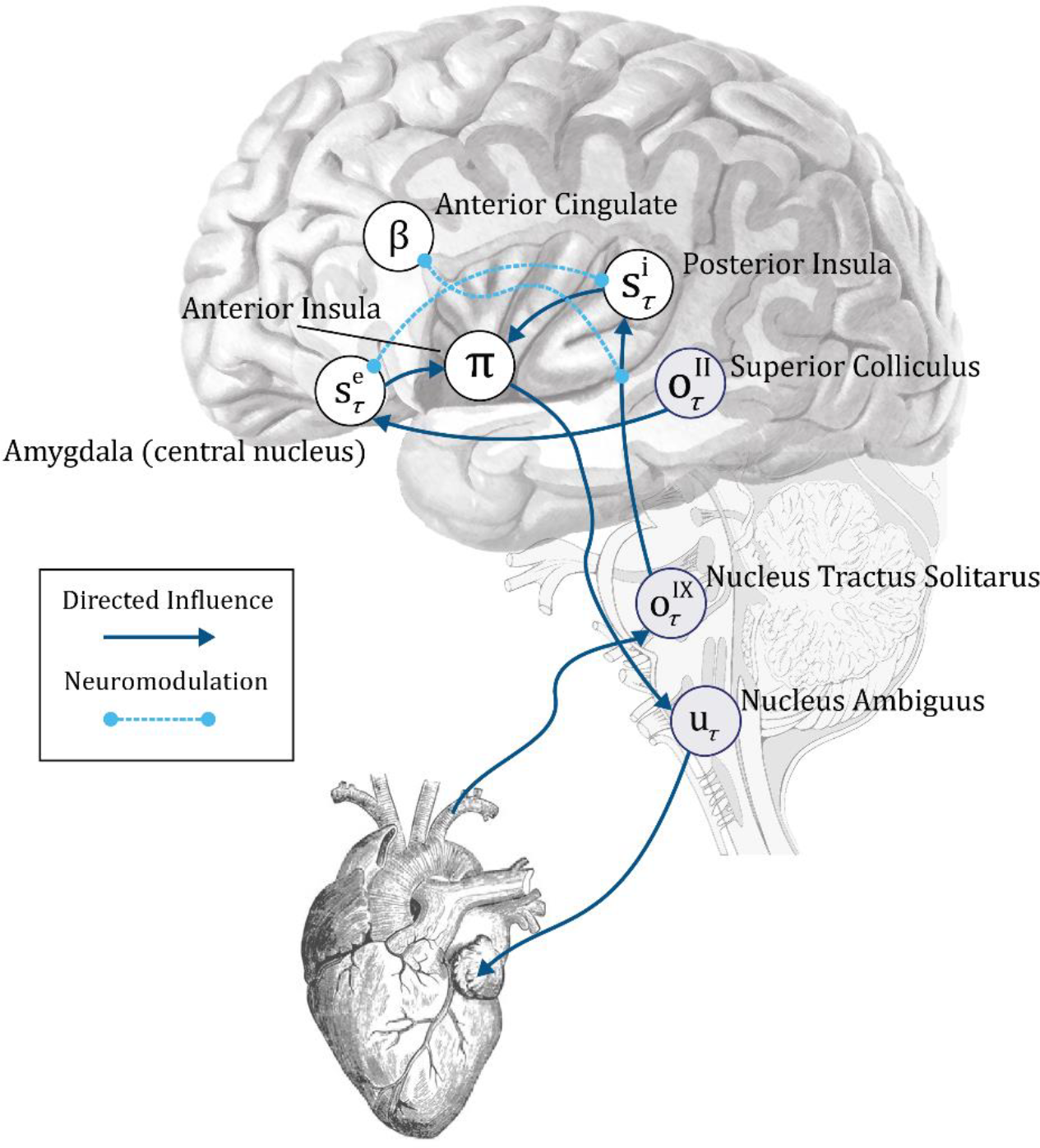
computational neuroanatomy of interoception. The schematic above shows the form of the neuronal message passing implied by active inference for the generative model depicted in Figure 2. We have related this to the anatomical networks that could implement these inferences. The sensory observations in our simulations are visual and interoceptive (cardiac). These sensations are carried by cranial nerves II and IX respectively. Cranial nerve II targets the superior colliculus in the midbrain. This structure sends short latency visual data to the amygdala, which is well placed to make inferences about emotionally salient stimuli. The amygdala additionally receives visual data from the ventral visual stream in the temporal lobe. Cranial nerve IX carries information from the carotid sinus baroreceptors to the nucleus tractus solitarus in the brainstem. This nucleus communicates with the posterior insula (via thalamic and PAG relays); the anterior cingulate monitors and controls the precision of this ascending visceral information via neuromodulation, possibly via feedback through noradrenergic pathways (not shown). The posterior insula and amygdala interact with one another but also project to the anterior insula. This targets the nucleus ambiguus (via brainstem relays such as the periaqueductal gray), which gives rise to the vagus (X) nerve. The vagus nerve targets neurons in the cardiac plexus that project to both the sinoatrial node and the atrioventricular node of the heart, slowing its rhythm. The nucleus tractus solitarus additionally participates in a reflex loop implicating the sympathetic control of the cardiac cycle, but this is omitted for simplicity. The functional anatomy suggested here implies the anterior insula might play a similar computational role in autonomic policy selection to the basal ganglia in selection of policies involving the skeletal muscles (Friston et al., 2018). Note that inscribed directed influences (blue arrows), are not assumed to be monosynaptic – for simplicity, many intermediary relay nodes have been omitted.

Within this scheme, we suggest that the rostral anterior cingulate (ACC) controls the precision of ascending visceral outcomes and inferred interoceptive states via neuromodulatory gain control (Fardo et al., 2017; Feldman and Friston, 2010). Further, interoceptive and exteroceptive state precisions (in our scheme) interact indirectly through global neuromodulatory influences, possibly through regulation of noradrenaline by the ACC (via descending influence on the locus coeruleus). Neurobiologically and functionally speaking, the AIC and ACC share similar profiles; both are densely populated with Von Economo neurons (VENs), which are well-suited for the long-range modulation of neural activity across the cortex (Allman et al., 2011), and also contain diverse populations of noradrenergic, dopaminergic, and opioidergic neurons. Both regions further share an integrative connectivity structure, with projections to both lower-level visceral-motor brainstem nuclei and higher-order regions implicated in decision-making, metacognition, and self-awareness, such as the ventromedial and dorsomedial prefrontal cortices (Allen et al., 2017, 2016a; Fleming and Dolan, 2012; Menon and Uddin, 2010; Ullsperger et al., 2010). However, the AIC is more densely interconnected with the PIC whereas the ACC is more closely related to uncertainty and decision-making. On this basis, we propose that whereas the AIC integrates the visceral and exteroceptive states required for the regulation of arousal policies, the ACC is likely to regulate the gain or precision of these interactions^4^.

What about metacognitive or reward-related interoceptive processes? Although here we do not model these higher-order functions, the model can be expanded to include the explicit representation of policy uncertainty and epistemic value as the mechanisms underlying metacognitive self-inference; i.e., the integrative self-model that combines exteroceptive and interoceptive predictions into a conscious schema (Allen and Tsakiris, 2019). In this case, we would expect that the VMPFC and DLPFC are likely to be engaged in inferences about variables (e.g., those derived from expected free energy such as epistemic and intrinsic value) that contextualize the inferences performed by the AIC and ACC over longer timescales (Friston et al., 2015, 2017a).

### Limitations and Future Directions

The model and simulations presented here represent a minimal proof-of-principle demonstrating how cyclic interactions of interoceptive and exteroceptive perception arise directly from the principles of active inference. Here, our primary goal was to move the literature beyond purely conceptual analyses of ‘interoceptive inference’, to provide a formal model sub-serving direct hypothesis testing. As such, we focus primarily on reproducing commonly reported phenomena, rather than empirical cross-validation or biological plausibility. While the model presented here does a reasonably good job of approximating the cardiac cycle, it should be clear that much work remains to be done if the model is to be used as a full generative model; e.g., of heart-brain interactions and/or physiological data such as HRV. We therefore anticipate a variety of fruitful applications. For example, the present MDP scheme could be expanded to include biologically realistic cardiac parameters, or to include other visceral modalities such as gastric or respiratory fluctuations. Similarly, the exteroceptive states modelled here could be adapted to a variety of experimental tasks to capture embodied influences on, for example, active spatial navigation (Kaplan and Friston, 2018; Lockmann et al., 2018; Lockmann and Tort, 2018), active reward learning (FitzGerald et al., 2015; Marshall et al., 2019), interaction between the cardiac cycle and ballistic saccades (Galvez-Pol et al., 2018; Mirza et al., 2016; Ohl et al., 2016), or metacognitive self-inference (Allen et al., 2016b; Friston et al., 2017b; Hauser et al., 2017a). These and other future directions will hopefully guide a newly embodied approach to computational psychiatry, enabling the detailed phenotyping of clinical populations in terms of aberrant interoceptive inference.

## Acknowledgements

MA is supported by a Lundbeckfonden Fellowship (R272-2017-4345), the AIAS-COFUND II fellowship programme that is supported by the Marie Skłodowska-Curie actions under the European Union’s Horizon 2020 (Grant agreement no 754513), and the Aarhus University Research Foundation. TP is supported by the Rosetrees Trust (Award Number 173346) TP. KJF is a Wellcome Principal Research Fellow (Ref: 088130/Z/09/Z). The authors further thank Maxwell Ramstead, Casper Hesp, and Francesca Fardo for fruitful discussions and inputs on the manuscript and modelling therein.

## Data and Code Availability

The underlying MDP scheme here is available as part of the open-access distribution of SPM12. A demonstration of the scheme can be accessed by typing >>DEM into the Matlab command prompt and selecting the **Interoception** demo from the graphical user interface that appears. Further, all the code required to generate the simulations and figures herein can be found at the following github page: https://github.com/embodied-computation-group/cardiac-active-inference.

## Footnotes

1 The term ‘belief’ here is used in the technical sense of a Bayesian belief, or probability distribution, typically considered to be sub-personal.

2 Every simulation started off with a weak prior over hidden states that the picture was not arousing – and a weaker prior in favor of the *relaxed* policy.

3 Sokolov (1963) described the orienting reflex as an ‘embodied’ mechanism for boosting to signal-to-noise and thus enhancing processing of the oddball stimulus. The reflex consists primarily of the rapid deployment of saccades to the oddball stimulus, freezing of the muscles of the head and neck so as to orient the visual organs towards the stimulus, and an immediate cardiac deceleration. In light of their inhibitory influence, the cardiac deceleration was thought to primarily reduce cortical noise; when coupled with the other bodily components of the response it was thought that effective overall signal would be maximized.

4 It is worth noting that this model may explain the widespread, seemingly unspecific activation profiles of these areas (Chang et al., 2013; Yarkoni et al., 2011), as the generative model specified here suggests both form part of an integrated hierarchical circuit by which interoceptive and exteroceptive states interact: e.g., either through the regulation of arousal policies or through the modulation of ascending viscerosensory precision.

## References

Allen M, Fardo F, Dietz MJ, Hillebrandt H, Friston KJ, Rees G, Roepstorff A. 2016a. Anterior insula coordinates hierarchical processing of tactile mismatch responses. NeuroImage 127:34–43. doi:10.1016/j.neuroimage.2015.11.030

Allen M, Frank D, Schwarzkopf DS, Fardo F, Winston JS, Hauser TU, Rees G. 2016b. Unexpected arousal modulates the influence of sensory noise on confidence. eLife 5:e18103. doi:10.7554/eLife.18103

Allen M, Friston KJ. 2018. From cognitivism to autopoiesis: towards a computational framework for the embodied mind. Synthese 195:2459–2482. doi:10.1007/s11229-016-1288-5

Allen M, Glen JC, Müllensiefen D, Schwarzkopf DS, Fardo F, Frank D, Callaghan MF, Rees G. 2017. Metacognitive ability correlates with hippocampal and prefrontal microstructure. NeuroImage 149:415–423. doi:10.1016/j.neuroimage.2017.02.008

Allen M, Tsakiris M. 2019. The body as first prior: Interoceptive predictive processing and the primacyThe Interoceptive Mind: From Homeostasis to Awareness. Great Clarendon Street, Oxford, OX2 6DP: Oxford University Press. pp. 27–45.

Allman JM, Tetreault NA, Hakeem AY, Manaye KF, Semendeferi K, Erwin JM, Park S, Goubert V, Hof PR. 2011. The von Economo neurons in the frontoinsular and anterior cingulate cortex. Ann N Y Acad Sci 1225:59–71. doi:10.1111/j.1749-6632.2011.06011.x

Anderson AK, Phelps EA. 2001. Lesions of the human amygdala impair enhanced perception of emotionally salient events. Nature 411:305–309. doi:10.1038/35077083

Apps MA, Tsakiris M. 2014. The free-energy self: a predictive coding account of self-recognition. Neurosci Biobehav Rev 41:85–97.

Attias H. 2003. Planning by probabilistic inference.Proc. of the 9th Int. Workshop on Artifical Intelligence and Statistics. Presented at the AISTATS.

Azevedo RT, Garfinkel SN, Critchley HD, Tsakiris M. 2017. Cardiac afferent activity modulates the expression of racial stereotypes. Nat Commun 8:13854. doi:10.1038/ncomms13854

Barto A, Mirolli M, Baldassarre G. 2013. Novelty or surprise? Front Psychol 4:907.

Birn RM. 2012. The role of physiological noise in resting-state functional connectivity. Neuroimage 62:864–870.

Boldt A, de Gardelle V, Yeung N. 2017. The impact of evidence reliability on sensitivity and bias in decision confidence. J Exp Psychol Hum Percept Perform 43:1520–1531. doi:10.1037/xhp0000404

Bonvallet M, Bloch V. 1961. Bulbar control of cortical arousal. Science 133:1133–1134.

Bonvallet M, Dell P, Hiebel G. 1954. Tonus sympathique et activité électrique corticale. Electroencephalogr Clin Neurophysiol 6:119–144.

Botvinick M, Toussaint M. 2012. Planning as inference. Trends Cogn Sci 16:485–488. doi:10.1016/j.tics.2012.08.006

Cechetto DF, Saper CB. 1987. Evidence for a viscerotopic sensory representation in the cortex and thalamus in the rat. J Comp Neurol 262:27–45. doi:10.1002/cne.902620104

Chang LJ, Yarkoni T, Khaw MW, Sanfey AG. 2013. Decoding the Role of the Insula in Human Cognition: Functional Parcellation and Large-Scale Reverse Inference. Cereb Cortex 23:739–749. doi:10.1093/cercor/bhs065

Chouchou F, Mauguière F, Vallayer O, Catenoix H, Isnard J, Montavont A, Jung J, Pichot V, Rheims S, Mazzola L. 2019. How the insula speaks to the heart: Cardiac responses to insular stimulation in humans. Hum Brain Mapp. doi:10.1002/hbm.24548

Cohen R, Lieb H, Rist F. 1980. Loudness judgments, evoked potentials, and reaction time to acoustic stimuli early and late in the cardiac cycle in chronic schizophrenics. Psychiatry Res 3:23–29.

Craig AD. 2002. How do you feel? Interoception: the sense of the physiological condition of the body. Nat Rev Neurosci 3:655–666. doi:10.1038/nrn894

Delfini LF, Campos JJ. 1972. Signal Detection and the “Cardiac Arousal Cycle.” Psychophysiology 9:484–491. doi:10.1111/j.1469-8986.1972.tb01801.x

Denève S, Jardri R. 2016. Circular inference: mistaken belief, misplaced trust. Curr Opin Behav Sci, Computational modeling 11:40–48. doi:10.1016/j.cobeha.2016.04.001

Eckberg Dwain L. 1997. Sympathovagal Balance. Circulation 96:3224–3232. doi:10.1161/01.CIR.96.9.3224

Edwards L, Ring C, McIntyre D, Winer JB, Martin U. 2009. Sensory detection thresholds are modulated across the cardiac cycle: Evidence that cutaneous sensibility is greatest for systolic stimulation. Psychophysiology 46:252–256. doi:10.1111/j.1469-8986.2008.00769.x

Elliott R. 1972. The significance of heart rate for behavior: A critique of Lacey’s hypothesis.

Fardo F, Auksztulewicz R, Allen M, Dietz MJ, Roepstorff A, Friston KJ. 2017. Expectation violation and attention to pain jointly modulate neural gain in somatosensory cortex. NeuroImage 153:109–121. doi:10.1016/j.neuroimage.2017.03.041

Feldman H, Friston K. 2010. Attention, Uncertainty, and Free-Energy. Front Hum Neurosci 4:215. doi:10.3389/fnhum.2010.00215

FitzGerald THB, Dolan RJ, Friston K. 2015. Dopamine, reward learning, and active inference. Front Comput Neurosci 9:136. doi:10.3389/fncom.2015.00136

Fleming SM, Dolan RJ. 2012. The neural basis of metacognitive ability. Philos Trans R Soc B Biol Sci 367:1338–1349. doi:10.1098/rstb.2011.0417

Fox MD, Raichle ME. 2007. Spontaneous fluctuations in brain activity observed with functional magnetic resonance imaging. Nat Rev Neurosci 8:700.

Fox MD, Snyder AZ, Vincent JL, Raichle ME. 2007. Intrinsic fluctuations within cortical systems account for intertrial variability in human behavior. Neuron 56:171–184.

Fox MD, Snyder AZ, Zacks JM, Raichle ME. 2006. Coherent spontaneous activity accounts for trial-to-trial variability in human evoked brain responses. Nat Neurosci 9:23.

Friston K, Rigoli F, Ognibene D, Mathys C, Fitzgerald T, Pezzulo G. 2015. Active inference and epistemic value. Cogn Neurosci 6:187–214.

Friston KJ, FitzGerald T, Rigoli F, Schwartenbeck P, Pezzulo G. 2017a. Active inference: a process theory. Neural Comput 29:1–49.

Friston KJ, Lin M, Frith CD, Pezzulo G, Hobson JA, Ondobaka S. 2017b. Active Inference, Curiosity and Insight. Neural Comput 29:2633–2683. doi:10.1162/neco_a_00999

Friston KJ, Parr T, de Vries B. 2017c. The graphical brain: Belief propagation and active inference. Netw Neurosci 1:381–414. doi:10.1162/NETN_a_00018

Friston KJ, Rosch R, Parr T, Price C, Bowman H. 2018. Deep temporal models and active inference. Neurosci Biobehav Rev 90:486–501. doi:10.1016/j.neubiorev.2018.04.004

Gallagher S, Allen M. 2018. Active inference, enactivism and the hermeneutics of social cognition. Synthese 195:2627–2648. doi:10.1007/s11229-016-1269-8

Galvez-Pol A, McConnell R, Kilner J. 2018. Active sampling during visual search is modulated by the cardiac cycle. bioRxiv 405902.

Garfinkel SN, Critchley HD. 2016. Threat and the Body: How the Heart Supports Fear Processing. Trends Cogn Sci 20:34–46. doi:10.1016/j.tics.2015.10.005

Garfinkel SN, Minati L, Gray MA, Seth AK, Dolan RJ, Critchley HD. 2014. Fear from the Heart: Sensitivity to Fear Stimuli Depends on Individual Heartbeats. J Neurosci 34:6573–6582. doi:10.1523/JNEUROSCI.3507-13.2014

Ghione S. 1996. Hypertension-associated hypalgesia: Evidence in experimental animals and humans, pathophysiological mechanisms, and potential clinical consequences. Hypertension 28:494–504.

Golanov EV, Yamamoto S, Reis DJ. 1994. Spontaneous waves of cerebral blood flow associated with a pattern of electrocortical activity. Am J Physiol-Regul Integr Comp Physiol 266:R204–R214. doi:10.1152/ajpregu.1994.266.1.R204

Graham FK, Clifton RK. 1966. Heart-rate change as a component of the orienting response. Psychol Bull 65:305–320. doi:10.1037/h0023258

Hassanpour MS, Yan L, Wang DJJ, Lapidus RC, Arevian AC, Simmons W. Kyle, Feusner Jamie D., Khalsa Sahib S. 2016. How the heart speaks to the brain: neural activity during cardiorespiratory interoceptive stimulation. Philos Trans R Soc B Biol Sci 371:20160017. doi:10.1098/rstb.2016.0017

Hauser TU, Allen M, Purg N, Moutoussis M, Rees G, Dolan RJ. 2017a. Noradrenaline blockade specifically enhances metacognitive performance. eLife 6. doi:10.7554/eLife.24901

Hauser TU, Allen M, Rees G, Dolan RJ. 2017b. Metacognitive impairments extend perceptual decision making weaknesses in compulsivity. Sci Rep 7:6614. doi:10.1038/s41598-017-06116-z

Heathers JAJ. 2012. Sympathovagal balance from heart rate variability: an obituary. Exp Physiol 97:556–556. doi:10.1113/expphysiol.2011.063867

Herrero JL, Khuvis S, Yeagle E, Cerf M, Mehta AD. 2017. Breathing above the brainstem: Volitional control and attentional modulation in humans. J Neurophysiol.

Itti L, Baldi P. 2009. Bayesian surprise attracts human attention. Vision Res 49:1295–1306.

Kaplan R, Friston KJ. 2018. Planning and navigation as active inference. Biol Cybern 112:323–343. doi:10.1007/s00422-018-0753-2

Karavaev AS, Kiselev AR, Runnova AE, Zhuravlev MO, Borovkova EI, Prokhorov MD, Ponomarenko VI, Pchelintseva SV, Efremova TY, Koronovskii AA, Hramov AE. 2018. Synchronization of infra-slow oscillations of brain potentials with respiration. Chaos Interdiscip J Nonlinear Sci 28:081102. doi:10.1063/1.5046758

Khalsa SS, Rudrauf D, Sandesara C, Olshansky B, Tranel D. 2009. Bolus isoproterenol infusions provide a reliable method for assessing interoceptive awareness. Int J Psychophysiol, Central and peripheral nervous system interactions: From mind to brain to body 72:34–45. doi:10.1016/j.ijpsycho.2008.08.010

Kiverstein J. 2018. Free Energy and the Self: An Ecological–Enactive Interpretation. Topoi. doi:10.1007/s11245-018-9561-5x

Koch E. 1932. Die irradiation der pressoreceptorischen kreislaufreflexe. J Mol Med 11:225–227.

Kunzendorf S, Klotzsche F, Akbal M, Villringer A, Ohl S, Gaebler M. 2019. Active information sampling varies across the cardiac cycle. Psychophysiology e13322.

Lacey BC, Lacey JI. 1978. Two-way communication between the heart and the brain: Significance of time within the cardiac cycle. Am Psychol 33:99.

Liddell BJ, Brown KJ, Kemp AH, Barton MJ, Das P, Peduto A, Gordon E, Williams LM. 2005. A direct brainstem–amygdala–cortical ‘alarm’ system for subliminal signals of fear. NeuroImage 24:235–243. doi:10.1016/j.neuroimage.2004.08.016

Limanowski J, Blankenburg F. 2013. Minimal self-models and the free energy principle. Front Hum Neurosci 7. doi:10.3389/fnhum.2013.00547

Lockmann ALV, Laplagne DA, Tort ABL. 2018. Olfactory bulb drives respiration-coupled beta oscillations in the rat hippocampus. Eur J Neurosci 48:2663–2673. doi:10.1111/ejn.13665

Lockmann ALV, Tort ABL. 2018. Nasal respiration entrains delta-frequency oscillations in the prefrontal cortex and hippocampus of rodents. Brain Struct Funct 223:1–3. doi:10.1007/s00429-017-1573-1

Malliani A, Pagani M, Lombardi F, Cerutti S. 1991. Cardiovascular neural regulation explored in the frequency domain. Circulation 84:482–492. doi:10.1161/01.CIR.84.2.482

Marshall AC, Gentsch A, Blum A-L, Broering C, Schütz-Bosbach S. 2019. I feel what I do: Relating interoceptive processes and reward-related behavior. NeuroImage 191:315–324. doi:10.1016/j.neuroimage.2019.02.032

Menon V, Uddin LQ. 2010. Saliency, switching, attention and control: a network model of insula function. Brain Struct Funct 214:655–667. doi:10.1007/s00429-010-0262-0

Mifflin SW, Felder RB. 1990. Synaptic mechanisms regulating cardiovascular afferent inputs to solitary tract nucleus. Am J Physiol-Heart Circ Physiol 259:H653–H661. doi:10.1152/ajpheart.1990.259.3.H653

Mirza MB, Adams RA, Mathys CD, Friston KJ. 2016. Scene Construction, Visual Foraging, and Active Inference. Front Comput Neurosci 10:56. doi:10.3389/fncom.2016.00056

Miura M, Reis DJ. 1972. The role of the solitary and paramedian reticular nuclei in mediating cardiovascular reflex responses from carotid baro- and chemoreceptors. J Physiol 223:525–548. doi:10.1113/jphysiol.1972.sp009861

Ohl S, Wohltat C, Kliegl R, Pollatos O, Engbert R. 2016. Microsaccades are coupled to heartbeat. J Neurosci 36:1237–1241.

Oppenheimer SM, Gelb A, Girvin JP, Hachinski VC. 1992. Cardiovascular effects of human insular cortex stimulation. Neurology 42:1727. doi:10.1212/WNL.42.9.1727

Oudeyer P-Y, Kaplan F. 2009. What is intrinsic motivation? A typology of computational approaches. Front Neurorobotics 1:6.

Owens AP, Allen M, Ondobaka S, Friston KJ. 2018. Interoceptive inference: From computational neuroscience to clinic. Neurosci Biobehav Rev 90:174–183. doi:10.1016/j.neubiorev.2018.04.017

Park H-D, Correia S, Ducorps A, Tallon-Baudry C. 2014. Spontaneous fluctuations in neural responses to heartbeats predict visual detection. Nat Neurosci 17:612–618. doi:10.1038/nn.3671

Park H-D, Tallon-Baudry C. 2014. The neural subjective frame: from bodily signals to perceptual consciousness. Philos Trans R Soc B Biol Sci 369:20130208.

Parr T, Friston KJ. 2018. The Computational Anatomy of Visual Neglect. Cereb Cortex 28:777–790. doi:10.1093/cercor/bhx316

Parr T, Friston KJ. 2017. Uncertainty, epistemics and active inference. J R Soc Interface 14:20170376. doi:10.1098/rsif.2017.0376

Parr T, Markovic D, Kiebel SJ, Friston KJ. 2019. Neuronal message passing using Mean-field, Bethe, and Marginal approximations. Sci Rep 9:1889. doi:10.1038/s41598-018-38246-3

Peters A, McEwen BS, Friston K. 2017. Uncertainty and stress: Why it causes diseases and how it is mastered by the brain. Prog Neurobiol 156:164–188. doi:10.1016/j.pneurobio.2017.05.004

Petzschner FH, Weber LAE, Gard T, Stephan KE. 2017. Computational Psychosomatics and Computational Psychiatry: Toward a Joint Framework for Differential Diagnosis. Biol Psychiatry, Computational Psychiatry 82:421–430. doi:10.1016/j.biopsych.2017.05.012

Powers AR, Mathys C, Corlett PR. 2017. Pavlovian conditioning–induced hallucinations result from overweighting of perceptual priors. Science 357:596–600. doi:10.1126/science.aan3458

Salomon R, Ronchi R, Dönz J, Bello-Ruiz J, Herbelin B, Martet R, Faivre N, Schaller K, Blanke O. 2016. The Insula Mediates Access to Awareness of Visual Stimuli Presented Synchronously to the Heartbeat. J Neurosci 36:5115–5127. doi:10.1523/JNEUROSCI.4262-15.2016

Sandman CA, McCanne TR, Kaiser DN, Diamond B. 1977. Heart rate and cardiac phase influences on visual perception. J Comp Physiol Psychol 91:189.

Saxon SA. 1970. Detection of near threshold signals during four phases of cardiac cycle. Ala J Med Sci 7:427.

Schmidhuber J. 2010. Formal theory of creativity, fun, and intrinsic motivation. IEEE Trans Auton Ment Dev 2:230–247.

Seth AK. 2013. Interoceptive inference, emotion, and the embodied self. Trends Cogn Sci 17:565–573. doi:10.1016/j.tics.2013.09.007

Seth AK, Friston KJ. 2016. Active interoceptive inference and the emotional brain. Philos Trans R Soc B Biol Sci 371:20160007. doi:10.1098/rstb.2016.0007

Seth AK, Tsakiris M. 2018. Being a Beast Machine: The Somatic Basis of Selfhood. Trends Cogn Sci 22:969–981. doi:10.1016/j.tics.2018.08.008

Sokolov EN. 1963. Higher nervous functions: The orienting reflex. Annu Rev Physiol 25:545–580.

Spence ML, Dux PE, Arnold DH. 2016. Computations underlying confidence in visual perception. J Exp Psychol Hum Percept Perform 42:671–682. doi:10.1037/xhp0000179

Strigo A, Craig AD. 2016. Interoception, homeostatic emotions and sympathovagal balance. Philos Trans R Soc B Biol Sci 371:20160010. doi:10.1098/rstb.2016.0010

Tort ABL, Brankačk J, Draguhn A. 2018a. Respiration-Entrained Brain Rhythms Are Global but Often Overlooked. Trends Neurosci 41:186–197. doi:10.1016/j.tins.2018.01.007

Tort ABL, Ponsel S, Jessberger J, Yanovsky Y, Brankačk J, Draguhn A. 2018b. Parallel detection of theta and respiration-coupled oscillations throughout the mouse brain. Sci Rep 8:6432–6432. doi:10.1038/s41598-018-24629-z

Ullsperger M, Harsay HA, Wessel JR, Ridderinkhof KR. 2010. Conscious perception of errors and its relation to the anterior insula. Brain Struct Funct 214:629–643. doi:10.1007/s00429-010-0261-1

Varga S, Heck DH. 2017. Rhythms of the body, rhythms of the brain: Respiration, neural oscillations, and embodied cognition. Conscious Cogn 56:77–90. doi:10.1016/j.concog.2017.09.008

Velden M, Juris M. 1975. Perceptual Performance as a Function of Intra-Cycle Cardiac Activity. Psychophysiology 12:685–692. doi:10.1111/j.1469-8986.1975.tb00075.x

Yarkoni T, Poldrack RA, Nichols TE, Van Essen DC, Wager TD. 2011. Large-scale automated synthesis of human functional neuroimaging data. Nat Methods 8:665–670. doi:10.1038/nmeth.1635

Zanatta P, Toffolo GM, Sartori E, Bet A, Baldanzi F, Agarwal N, Golanov E. 2013. The human brain pacemaker: Synchronized infra-slow neurovascular coupling in patients undergoing non-pulsatile cardiopulmonary bypass. NeuroImage 72:10–19. doi:10.1016/j.neuroimage.2013.01.033

Zelano C, Jiang H, Zhou G, Arora N, Schuele S, Rosenow J, Gottfried JA. 2016. Nasal Respiration Entrains Human Limbic Oscillations and Modulates Cognitive Function. J Neurosci 36:12448–12467. doi:10.1523/JNEUROSCI.2586-16.2016

